# Deciphering complex mechanisms of resistance and loss of potency through coupled molecular dynamics and machine learning

**DOI:** 10.1101/2020.06.08.139105

**Authors:** Florian Leidner, Nese Kurt Yilmaz, Celia A. Schiffer

**Affiliations:** Department of Biochemistry and Molecular Pharmacology, University of Massachusetts Medical School, Worcester, Massachusetts 01605, United States

## Abstract

Drug resistance threatens many critical therapeutics through mutations in the drug target. The molecular mechanisms by which combinations of mutations, especially involving those distal from the active site, alter drug binding to confer resistance are poorly understood and thus difficult to counteract. A machine learning strategy was developed that couples parallel molecular dynamics simulations and experimental potency to identify specific conserved mechanisms underlying resistance. A series of 28 HIV-1 protease variants with 0-24 substitutions each were used as a rigorous model of this strategy. Many of the mutations were distal from the active site and the potency of variants to a drug (darunavir) varied from low pM to near µM. With features extracted from the simulations, elastic network machine learning was applied to correlate physical interactions with loss of potency and succeeded to within 1 kcal/mol of experimental affinity for both the training and test sets, outperforming MM/GBSA calculations. Feature reduction resulted in a model with 4 specific features that describe interactions critical for potency for all 28 variants. These predictive features, that specifically vary with potency, occur throughout the enzyme and would not have been identified without dynamics and machine learning. This strategy thus captures the conserved dynamic mechanisms by which complex combinations of mutations confer resistance and identifies critical features that serve as bellwethers of loss of inhibitor potency. Machine learning models leveraging molecular dynamics can thus elucidate mechanisms of drug resistance that confer loss of affinity and will serve as predictive tools in future drug design.

## Introduction

Drug resistance is ubiquitous in both infectious diseases and oncology, impeding the success of drug therapy and severely impacting human health. ^1, 2^ Mechanisms that confer drug resistance are manifold and include reduced cellular uptake, degradation, and changes in target expression. ^1^ Often drug resistance is caused by mutations that disrupt drug–target interactions while preserving the target’s function in the presence of drug. Drug resistance due to loss of drug– target interactions is particularly prevalent in viruses and low complexity organisms that do not possess the genomic capacity to encode for auxiliary resistance genes but occurs in all quickly evolving drug targets. ^3-5^

HIV-1 is one of the most rapidly evolving viruses and treatment of HIV-1 infections has historically been severely affected by drug resistance. ^6, 7^ The fast rate of mutation and abundance of experimental data make HIV-1 protease an ideal model system to elucidate the molecular mechanisms by which mutations cause resistance. Structural studies on HIV-1 protease and drug resistance have led to the development of the substrate envelope hypothesis, which postulates that competitive active site inhibitors can avoid resistance by maximizing shape similarity with the native substrates. ^8^ The success of this strategy has been demonstrated not only in HIV-1 protease but also in other rapidly evolving viral targets. ^9, 10^ The most recently FDA approved HIV-1 protease inhibitor darunavir (DRV) adheres to this strategy, resulting in a remarkably high barrier to resistance. Resistance to DRV still emerges through accumulation of multiple mutations both proximal and distal from the active site. ^11-14^ Elucidating the resistance mechanism for very potent inhibitors, such as the HIV-1 protease inhibitor DRV, provides insights into how amino acid substitutions alter the structure, dynamics and function of a therapeutic target. ^15, 16^ However the molecular mechanisms by which mutations, in particular those distal from the active site, confer resistance remains elusive.

A common assumption is that these distal mutations are required to retain structural stability and enzymatic activity by compensating for otherwise deleterious active site mutations. ^17, 18^ However in-depth studies of highly resistant drug resistant protease variants indicate that distal mutations can also alter drug–target interactions through propagating dynamic alterations within the network of intra-protein interactions, as proposed by the “network hypothesis”. ^19, 20^ Consistent with this, NMR studies indicate that drug resistant variants show distinct changes in protein dynamics. ^21^ Our initial regression models where we examined one physical feature at a time also indicated that dynamics were likely the key in resistance ^22, 23^. We previously characterized a series of increasingly drug resistant protease variants from viral passaging experiments and through co-crystal structures, enzyme inhibition and molecular dynamics (MD) simulations demonstrated that variants with mutations distal from the active site weakened interactions with the inhibitor through changes in structure and dynamics. ^24^

Although enzyme–inhibitor dynamics are known to be critical in resistance especially due to mutations distal from the active site, the molecular mechanisms involved are not clear. Resistance may be conferred through dynamic changes specific to a given variant, or through conserved mechanisms that can be captured by a comprehensive analysis of an enzyme– inhibitor system. Identifying such conserved mechanisms would enable predicting the cause of drug resistance and designing more robust inhibitors to avoid resistance. Here we systematically probe the ensemble dynamics of HIV-1 protease variants with increasing levels of resistance, to elucidate the molecular mechanisms of drug resistance. Machine learning was used to construct a model correlating physical interactions at the molecular level with the loss of inhibitor potency, based on features calculated from MD simulations. We found that as few as four physical features were sufficient to describe the observed changes in binding affinity across a broad range of HIV-1 protease variants. These physical features include direct interactions between protease and inhibitor, as well as intra-molecular contacts between distal protein residues which likely anchor the protease in the closed, inhibited conformation. The accuracy of the resulting model exceeds that of endpoint free energy calculations, which fail to account for the impact of distal changes. The predictive accuracy of the model was further validated on a separate test set of protease variants not used in building the model. Machine learning models identifying and leveraging such “bellwether” interactions can thus strongly correlate with inhibitor potency and serve as predictive tools in drug design.

## Results

A set of 28 HIV-1 protease variants was chosen to cover a wide range of darunavir (DRV) susceptibility and resistance, over six orders of magnitude in potency. The variants included up to 24 mutations each, with substitutions in 48 of the 99 residues within the enzyme. (Fig 1; Fig. S1). The number and distribution of the amino acid substitutions are in excellent agreement with protease variants observed in clinical isolates. ^25^ The spatial distribution of mutations (Fig. 1A) highlights the remarkable genomic plasticity of HIV-1 protease; contiguous regions without changes are found only surrounding the catalytic site and at the terminal dimerization motif. As HIV-1 protease is a homodimer (residues on either of the two monomers are indicated by a subscript A or B below) each mutation has a double impact. Only a few sites had ≥3 substitutions (Fig. 1B) including the polymorphic residues 63 and 89 and three resistance-associated sites 10, 54 and 82. ^26, 27^ Most substitutions in these resistant variants are distal from the inhibitor binding site.

**Figure 1.**
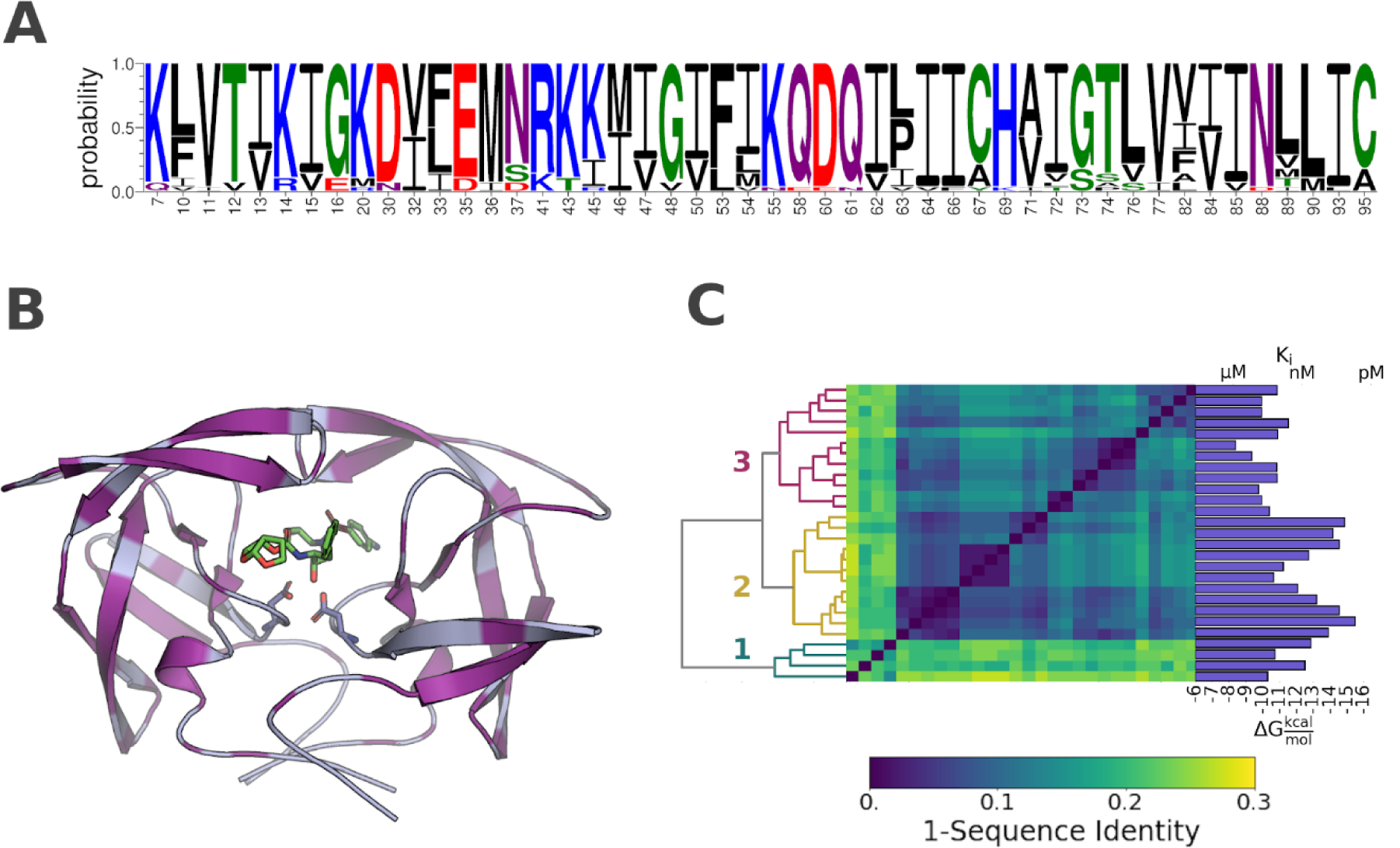
Amino acid sequence variation and distribution in the HIV-1 protease variants: **A)** Sequence logo showing all residues with one or more amino acid substitution. **B)** Structure of HIV-1 in complex with darunavir (PDB ID: 6dgx). Protease shown as light blue cartoon. Sites of mutations highlighted in purple. Inhibitor and the catalytic aspartic acids displayed as sticks **C)** Hierarchical clustering of the HIV-1 protease variants according to sequence identity. Dendrogram shown on the left, dissimilarity matrix in the center, and darunavir potency for each variant shown on the right.

The protease variants clustered into 3 major subgroups based on amino acid sequence. (Fig 1C, Table S1) Cluster 3 included all variants obtained from viral passaging experiments with DRV and DRV analogs ^14^. Cluster 2 included variants with less than 5 amino acid substitutions. Finally, cluster 1 included all protease variants obtained from clinical isolates. Cluster 1 was by far the most diverse, having a mean sequence identity of 80% with the remaining protease variants ^11, 28^. Despite the larger mutational load, the clinical isolates were not more resistant to DRV than the variants obtained from viral passaging. In fact, the number of substitutions was only weakly correlated with DRV potency. Variants with less than 5 mutations were significantly more susceptible to DRV; however, for the clinical isolates and for variants obtained from viral passaging experiments, no simple relationship could be established between the number of substitutions and susceptibility to DRV.

### Feature Selection: Determination of Specific Features

To characterize changes in interactions and dynamics across the HIV-1 protease variants, 100 ns simulation of all 28 variants were performed in triplicates. A total of 2858 features were calculated from molecular dynamics simulations for each complex. These features included van der Waals (vdW) interactions, hydrogen bonds, torsion angle entropy and root mean squared fluctuations. As molecular features also contain information about amino acid sequence identity, a critical test was to see whether the effect of any given molecular feature could equally be explained by the mere presence of a particular sequence mutation. Molecular features that substitute for sequence information decrease the specificity of the model, as sequence alone does not ascertain which specific physical changes occur at the molecular level. Such features would also reduce the generalizability of the model, as only a limited number of amino acid combinations can be included in building the model.

For regression analysis a parsimonious set of features is generally desirable, to improve the model accuracy and interpretability. ^29^ Therefore only the subset of features that strongly impact potency were selected. Features were selected according to three criteria: Accuracy, Stability and Specificity. Accuracy requires the features to be strongly correlated with the observed binding affinity. Stability requires that the weight of a feature be unaffected by relatively minor perturbations in the data. Finally, specificity requires that the feature inform on the relationship between changes in molecular interactions and changes in potency. Accuracy and stability were achieved by using regularized regression model on a subset (66%) of the training set and repeating this process on random permutations of the entire training set of variants. Features were defined to be specific when they provided more information on inhibitor potency than an alternative regression model trained on amino acid sequence information alone.

Elastic net regression was used to identify which among the 2858 features best correlated with the observed potency. The model was fit to two thirds of the training data while one third was left for model evaluation. To ensure convergence of the resulting coefficients, this process was repeated 100 times with random permutations of the dataset. 59 features with non-zero coefficients (p<0.05; 1 sample t-test) were identified. When sorted, the mean absolute coefficients exhibited an exponential decay with the knee of the curve located at 0.02. (Fig. 2A) Only 9 of the selected features had coefficients above 0.2, which are anticipated to be the most correlated with potency.

**Figure 2.**
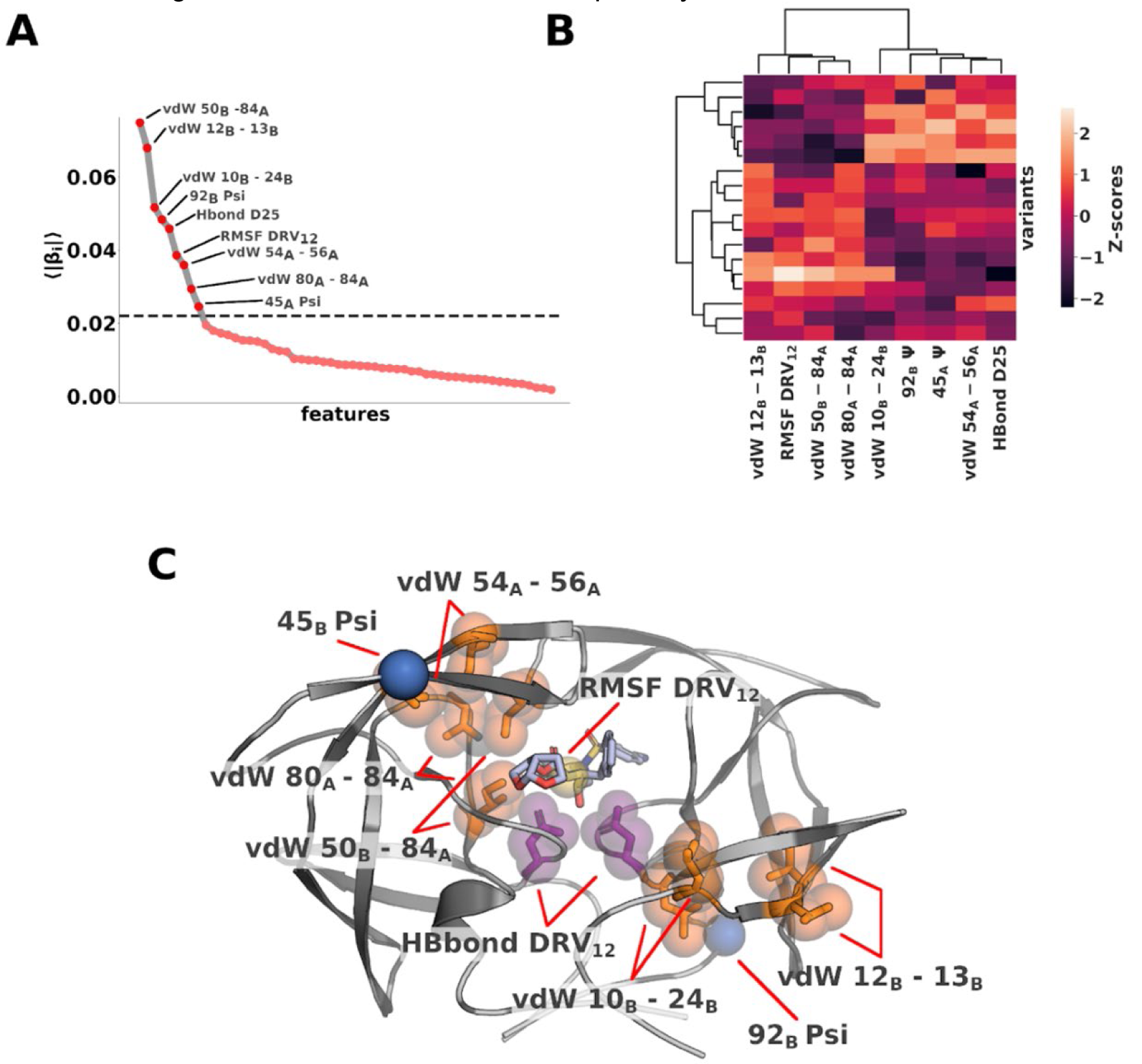
Molecular features indicative of inhibitor potency **A)** Absolute coefficients averaged over 100 rounds of shuffle-split cross validation. 9 features have coefficients larger than 0.02 (dashed horizontal line) **B)** Hierarchical clustering of the top 9 features according to coefficients **C)** Features with top 9 coefficients mapped onto the structure of protease–DRV complex. vdW interactions shown in orange, RMSF in yellow, torsion angle in blue and hydrogen bond in purple.

The 9 remaining features included 5 intra-protein vdW interactions, the root-mean-squared fluctuations of one inhibitor nitrogen atom (RMSF DRV12), two torsion angles (92_B_ Ψ and 45_B_ Ψ), and the hydrogen bond between the catalytic residues and inhibitor (Hbond D25). Except for RMSF DRV12, 92B Ψ and Hbond D25, all features involved at least one variable residue. The 9 selected features could be separated into two distinct clusters based on their correlation with one another indicating the complexity of their interdependence. (Fig. 2B) Physically, when mapped onto the protein structure, the residues involved in the interactions formed a contiguous region bisecting the protease dimer diagonally from the “flap”, through the active site, to the outer loops of the dimer. (Fig. 2C) Interestingly, on the protein structure these features have an asymmetric arrangement. HIV-1 protease is a homodimer; however, asymmetry is introduced when a non-symmetric inhibitor is bound. To establish whether the observed asymmetry is a determining factor or the result of a random selection process, all symmetry related features were swapped (e.g. 92_B_ Ψ → 92_A_ Ψ). DRV12 and H-Bond D25 were kept the same as there is no symmetry-related counterpart. The symmetry swapped model performed significantly worse than the original model, confirming that the features are specific to one protein chain. Most of the specific features involved hydrophobic residues. Rearrangements in the hydrophobic core, or hydrophobic sliding, of the protease has previously been proposed by us as a potential mechanism by which distal mutations alter inhibitor interactions to confer resistance. ^30, 31^ Our findings not only corroborate these results, but the selected features also indicate the key interactions associated with such changes that translate to alterations in potency.

### Predictive Models of Potency

The best model to predict potency using the 9 selected features with the least possible parameters was then determined. The features retained significant collinearity, implying potential redundancy or interdependency. (Fig. 2B) All 511 feature combinations were evaluated to converge on the final model. After discarding all combinations with a relative likelihood of minimizing the information loss < 0.05 (See Methods, Eq. 3 and Eq. 4), the final model was chosen by minimizing the number of parameters and maximizing the coefficient of determination (r^2^). This model included only 4 features (*vdW 12B-13B, vdW 10B-24B, vdW 50B-84A* and *HBond D25*). Using 5-fold cross validation, the root mean squared error (RMSE) of the model with these 4 features on the training set was 0.9 kcal/mol, Pearson correlation was 0.9 and Spearman correlation 0.8. (Fig. 3) On the independent test set, the RMSE was 1.3 kcal/mol and correlation coefficients dropped to 0.6 and 0.5. (Fig. 3) The performance on the test set captures the predictive ability and generalizability of the model. Including more than 4 parameters did not improve the fit. In fact, including all 9 parameters in the model significantly increased the variance of the predictions and only had a marginal effect on the r^2^: RMSE on the training and test sets decreased to 1.2 kcal/mol and 1.5 kcal/mol respectively, with worse p-values (>0.05). Thus, the model with 4 parameters performed the best. This level of accuracy was reassuring of physical reasonableness of the model since the parameters were selected using the full training set and the model could translate predictive power to naïve data.

**Figure 3.**
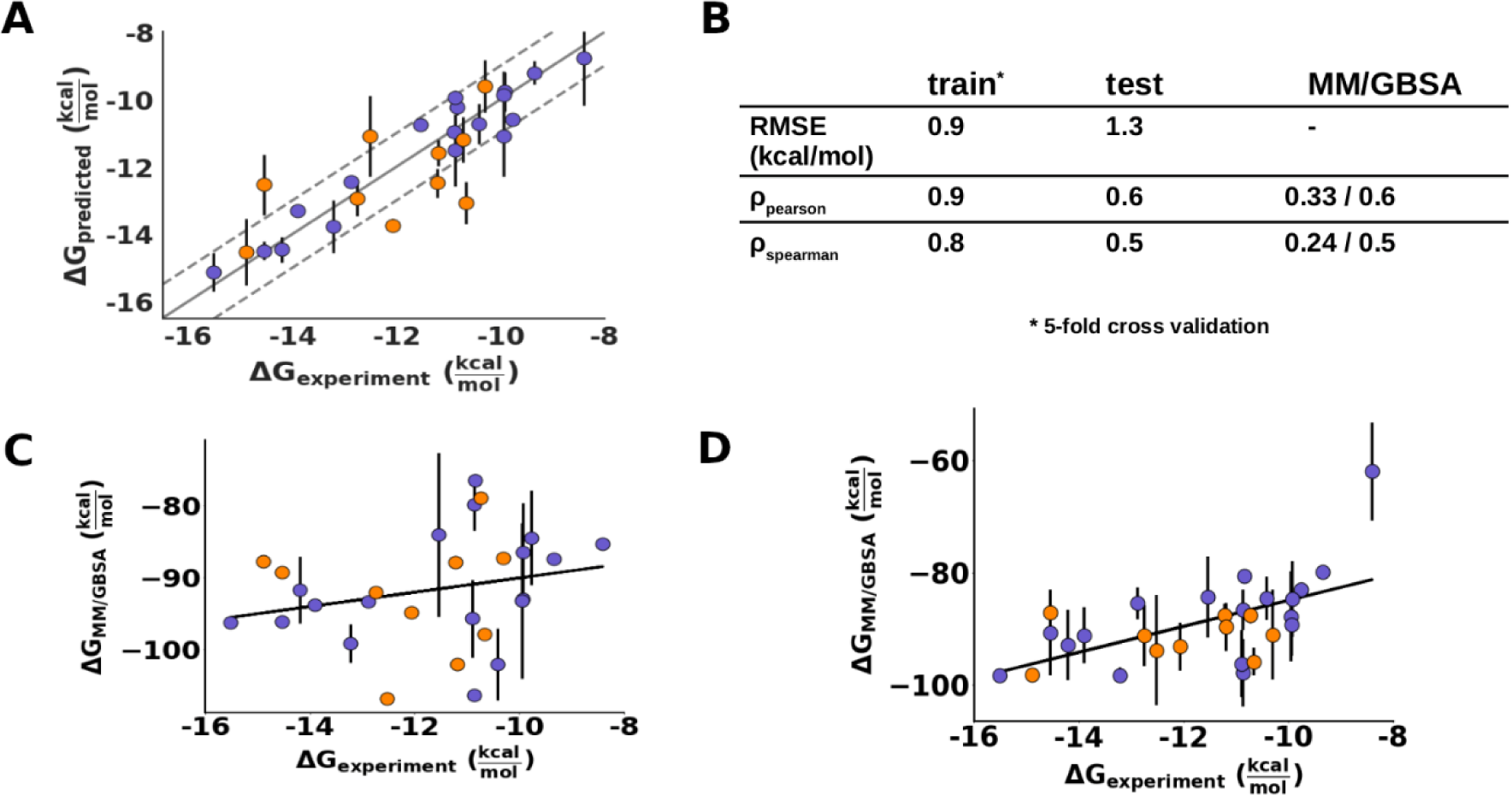
**A)** Evaluation of the regression model on training (purple dots) and test (orange dots) sets for prediction of DRV potency with only 4 features. Error bars indicate the 95% confidence interval. Training set metrics were calculated using 5-fold cross validation. **B)** Fit with experimental data. The correlation values for MM/GBSA correspond to the single frame and full trajectory results. **C)** MM/GBSA binding free energy of DRV against training and test sets averaging 3 snapshots from the trajectory. **D)** MM/GBSA binding free energy of DRV against training and test sets averaging the entire trajectory. Error bars correspond to the 95% confidence interval estimated from the 3 simulation replicates.

To compare the performance of our model with methods commonly used to evaluate ligand affinity, the binding free energy of DRV was calculated using MM/GBSA. ^32, 33^ Endpoint methods such as MM/GBSA are agnostic to dynamics and merely compare the free energy of the bound and unbound states. This method has been previously used to evaluate the impact of amino acid substitutions in HIV-1 protease. ^34^ MM/GBSA is historically performed on a single conformation, mostly with the inhibitor docked to a protein crystal structure. For a single structure, MM/GBSA scores did not correlate with the binding potency (ρ_Pearson_: 0.33; p: 0.001). To attempt a comparison that also includes dynamics, the binding free energy was calculated in 1 ns intervals during every MD simulation (3 replicates of 100 ns), and averaged over the resulting 300 calculations per protease variant. This resulted in a far better (ρ_Pearson_: 0.58; p: 0.001) correlation between the experimental potency and the binding free energy calculated from MM/GBSA, validating that dynamics is key to characterizing potency. (Fig. 4) Previous studies that combined MD and MM/GBSA typically used much shorter simulations (4–40 ns) and had low accuracy to fitting the data. ^35, 36^ The most resistant variant was a notable outlier in the MM/GBSA calculations even with dynamics included. We have characterized this variant extensively in the past and found its conformational dynamics to deviate significantly from those of other protease variants. ^24^ Because this highly resistant variant samples a broader distribution of the bound states, the average free energy deviates significantly from the other estimates. Overall, even though the free energy calculations performed better when averaged over a set of snapshots from the MD simulations, the performance was still considerably worse than our regression model.

**Figure 4.**
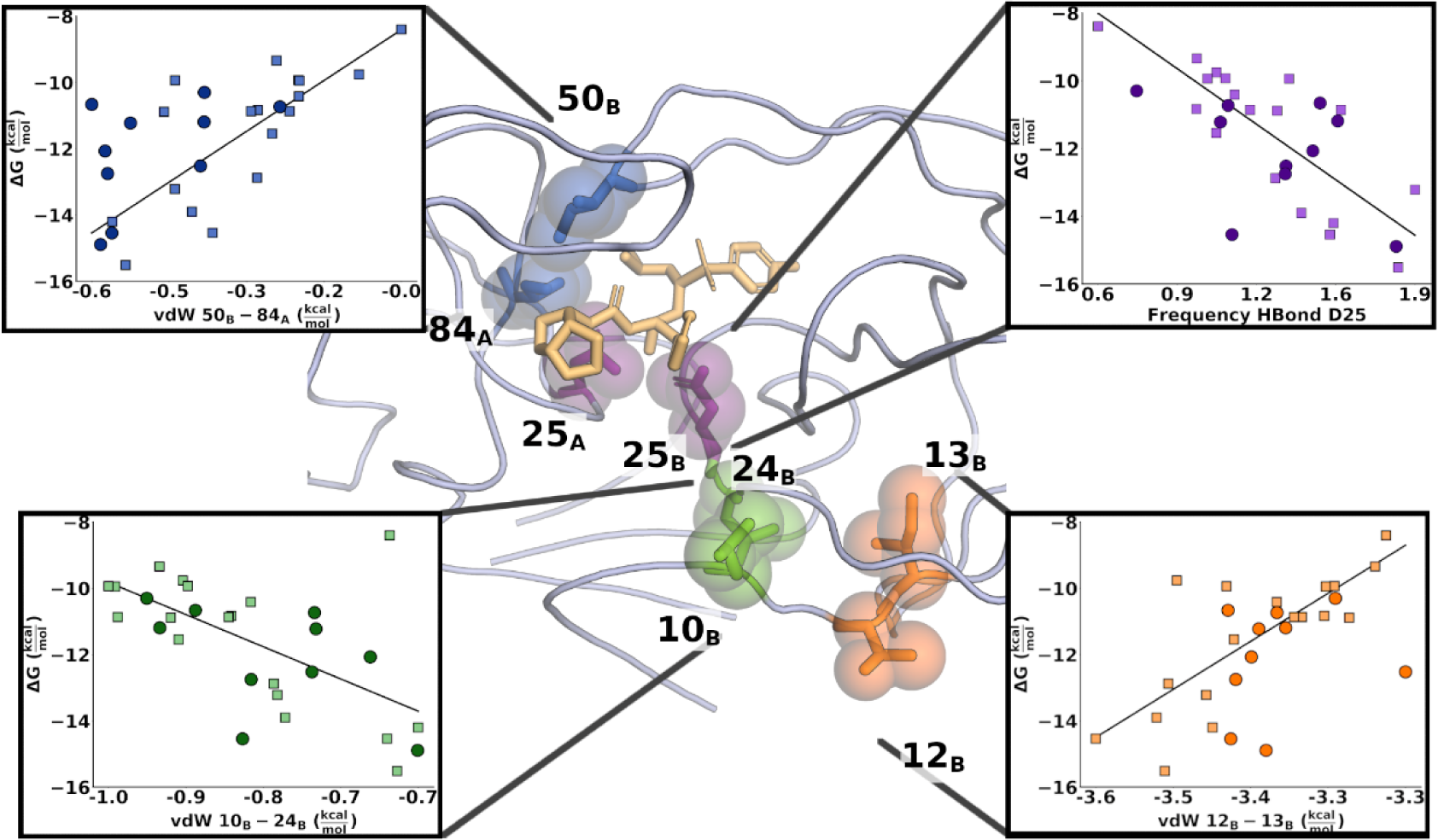
Signature interactions mapped onto the structure of HIV-1 protease. DRV shown in beige sticks. Insets show linear regression curves for each feature, with training set datapoints shown as squares and test set datapoints shown as circles. Dark blue: vdW interactions between residues 50_B_ and 84_A_; Purple: the hydrogen bond between the catalytic residue D25 and DRV; Green: vdW interactions between residues 10_B_ and 24_B_; and Orange: vdW interactions between residues 12_B_ and 13_B_.

### Molecular Indicators of Potency

As the potency predicted from our model was in excellent agreement with the experimental data and retained accuracy against the test set (Fig 3), the 4 specific features used in the predictive model were investigated in detail. These features indicate physical properties and interactions implicated in molecular mechanism of resistance. The weight of the feature in the regression model are provided in Table 1. The feature, that showed the strongest relationship with potency was the vdW interaction between the distal protease residues 12_B_-13_B_. In all but 2 variants, residue 12 is a threonine whereas residue 13 is either an isoleucine (WT) or a valine.(Figure S1) At first this result appears confounding, as these two residues neither make direct contact with the inhibitor nor face each other. However, a decrease in short-range interactions between residues 12 and 13 correlates strongly with decreased DRV binding affinity, consistently in both training and test sets. (Fig. 4) As these two residues are adjacent to each other, the loss of interactions can be explained by either a mutation or changes in relative sidechain orientation. If the effect was purely an indicator of amino acid identity, the feature would have been eliminated during feature selection, thus the orientation of the side chains of residues 12 and 13 are predictive for DRV affinity. Residue 13 is a common polymorphic mutation in other (non-B) HIV subtypes and the preexistence of 13V in treatment-naïve viral populations can lower the genetic barrier to resistance. ^37^ To identify whether the side chain orientation of these two residues relates to changes in specific molecular interactions, all features were regressed against vdW 12_B_-13_B_. Features that showed a strong linear relationship with vdW 12_B_-13_B_ (r^2^ ≥ 0.65) were in the immediate physical vicinity on the protein structure and mostly encoded for vdW interactions between 13_B_ and other hydrophobic residues, suggesting that vdW 12_B_-13_B_ acts as an indicator of the packing in the hydrophobic core, which as we and others previously suggested is reorganized due to resistance. ^30, 31, 38^

**Table 1.** Ordinary least squares model (Eq. 5) of DRV binding as a function of intra- and intermolecular interactions. β are the coefficients, σ the standard deviation and p the probability of H_0_: β=0T.

| | $\beta$ | $\sigma$ | $p$ |
| --- | --- | --- | --- |
| **intercept** | -11.50 | 0.16 | 0.0 |
| **vdW 10<sub>B</sub>-24<sub>B</sub>** | -0.58 | 0.17 | 0.005 |
| **vdW 12<sub>B</sub>-13<sub>B</sub>** | 0.73 | 0.20 | 0.003 |
| **vdW 50<sub>B</sub>-84<sub>A</sub>** | 0.59 | 0.23 | 0.02 |
| **HBond D25** | -0.55 | 0.24 | 0.04 |

A similar trend was observed for the vdW interactions between residues 10_B_ and 24_B_. Changes in these interactions were coupled with a weakening of the backbone hydrogen bond between residues 9_B_ and 24_B_, which in turn correlated with changes in both vdW and hydrogen bond interactions in the active site. Of note is that vdW 10_B_-24_B_ and vdW 12_B_-13_B_ were associated with changes in two different regions of the enzyme, suggesting that these features are indicative of distinct rearrangements.

The third feature involved vdW interactions within the active site, between residues 50_B_-84_A_. Both residues are prominent sites of drug resistance mutations (I50V/L and I84V) which can contribute significantly to weakening inhibitor interactions. Residue 50 is located on the tip of the protease “flap”, the anti-parallel beta sheet acting as a gate to the active site. Sidechain of residue 50_B_ contacts the side chain of residue 84_A_, across the dimer interface. Decreased vdW interactions between these two residues are associated with a loss of potency and indicate weakening of the inhibited, closed form of the enzyme and the slight opening of the “B” monomer flap. Changes in vdW 50_B_-84_A_ interactions were only weakly related to other changes throughout the enzyme. However, the interactions may also be affected by other mutations, because the most significant loss in 50_B_-84_A_ interactions was observed in variants that had only the I84V mutation, whereas variants that had mutations at both I50 and I84 mutations showed only a moderate change in interactions. Thus, distal mutations can propagate effects to the active site to weaken the critical residue 50-84 interactions that are indicative of closed flap (or inhibited) conformation.

The final interaction included in the model was the hydrogen bond between the catalytic aspartic acids and DRV. Loss of this pivotal hydrogen bond was associated with a decrease in inhibitor binding. Decreased interactions with the catalytic aspartates have in the past been implicated to play a role in resistance against HIV-1 protease inhibitors. ^39, 40^ Our results reinforce these findings and display a linear relationship between DRV binding and the stability of the D25 hydrogen bond. Overall we identified select and specific interactions indicative of concerted rearrangements in the hydrophobic core of the enzyme (vdW 10_B_-24_B_ and vdW 12_B_-13_B_), changes in the active site (vdW 50_B_-84_A_), and interactions with the bound inhibitor (HBond D25); these can be seen as bellwethers of loss of potency, in this case, due to drug resistance.

## Discussion

Elucidating the molecular mechanisms of resistance is critical for the design of inhibitors that can avoid resistance and retain potency against rapidly evolving disease targets. Here we demonstrate that combining machine learning with molecular dynamics and experimental affinity enables generating predictive models that identify molecular interactions indicative of decreased inhibitor affinity due to resistance over a wide range of potency. Molecular dynamics simulations and analysis of structural and dynamic features can quantify changes in molecular interactions; however, selecting a set of parameters that describe and predict inhibitor potency remains challenging. Thus, we developed a machine learning protocol that generated a predictive model of potency from merely 4 easily interpretable physical features. The predicted change in potency is in very good agreement with experimental values, outperforming MM/GBSA calculations. Furthermore, the model retained accuracy against a test set of variants not used in the training process, demonstrating the ability to generalize the model to unseen data. Many protease variants had vastly different (Fig. S1) combinations of mutations yet resistant variants ultimately converged to the same molecular feature phenotypes. Critically, the predictive accuracy of our model over these variants indicates that drug resistance is mediated through conserved and specific mechanisms that are independent of the particular sites of mutations and can be characterized by select few molecular interactions. Due to the feature selection and design of the machine learning protocol, this correlation is not a mere indicator of changes in amino acid sequence, but rather reflects alterations in molecular interactions and dynamics of the enzyme inhibitor complex. Perhaps more importantly, these conserved interactions can be used to predict potency of novel inhibitors with improved resistance profiles.

Proteins, in particular enzymes, are dynamic molecules recognizing substrates, processing and releasing products. When an enzyme is a quickly evolving drug target, resistance emerges such that the balance of recognition favors the dynamics of substrate binding over inhibition. Thus, to elucidate the underlying mechanisms of drug resistance, assessing molecular dynamics is essential. Mutations in resistant variants of enzymes that are targeted with therapeutics occur throughout the structure both within the active site and at distal positions. Primary resistance mutations at the active site can effectively be explained by the substrate envelope model, where the mutation preferentially weakens inhibitor binding. ^8^ However, explaining how distal mutations confer resistance is more challenging. In our study of HIV-1 protease we have demonstrated how the dynamics of the enzyme is impacted by a combination of distal and active site mutations ^19, 22, 23^, and here comprehensive analysis with machine learning identified conserved mechanisms by which resistance occurs.

Distal mutations cause resistance in many therapeutic target enzymes. These include kinases such as BCR-ABL with at least 19 mutations implicated in resistance ^41^, and EGFR with 13 sites of resistance characterized ^42-46^. In antibacterials, sequence changes can cause potency loss, such as variants of dihydrofolate reductase resistant to the widely used drug trimethoprim ^47-49^. Variations in sequence including those distal from the active site can also be critical for development of pan-viral inhibitors against diseases such as those caused by flaviviruses (Dengue, Zika and Yellow Fever), and coronaviruses such as SARS-CoV, MERS-CoV and SARS-CoV-2. In addition, variations distal from the active site are key when developing specific inhibitors for a certain enzyme in a family, with identical active sites. Thus, methods to assess structural and dynamic features impacted by not only the active site but also distal mutations are required in many drug design applications. Combining machine learning with parallel molecular dynamics and experimental data to identify key bellwethers of potency will likely become a powerful strategy in drug design.

## Materials and Methods

The code to calculate molecular features, the full set of features and the regression analysis are made available on Github (https://github.com/SchifferLab/ROBUST).

### Data Curation

#### HIV-1 protease variants and experimental inhibitor potency

Drug resistant variants of HIV-1 protease were obtained from viral passaging experiments with DRV and closely related analogs. ^50^ These variants were supplemented with variants bearing known primary active site mutations. The crystal structures of wild type protease, variants with primary resistance mutations, and highly drug resistant variants bound to DRV had been determined previously. ^24, 51^ The experimental K_i_ measurements were converted into free energy values using R×T×log(K_i_), where R is the gas constant in kcal*K^-1^*mol^-1^. The temperature was set to 300 K. To supplement the set of in-house variants, a set of resistant variants from the literature was curated. The set was compiled to resemble the spread in free energy values observed in the in-house dataset. For this test set, inhibition constant reported in the primary reference were used. (Table S1)

#### Structure preparation

Protein–inhibitor complex structures were retrieved from the PDB. ^52^ When DRV was found in multiple conformations, we chose the conformation with the highest mean occupancy or the first conformation listed in PDB file in cases of equal occupancy. In one crystal structure (3ucb; Table1) a second molecule of DRV was co-crystallized outside of the active site and was removed for this analysis. All variants from viral passaging (Source: ^50^, Table1) that lacked experimental structures were modeled using Prime. ^53, 54^

Structures were prepared using the *Schrödinger Protein Preparation Wizard*. ^55^ The protonation states were calculated using Propka. Protonation of the two catalytic aspartate residues, which exist primarily in the monoprotonated form, was examined and, if necessary, adjusted using the pKa values predicted by Propka to determine which of the two aspartic acids was protonated. ^56, 57^ If necessary, the chain ID of the two monomers were exchanged to ensure uniformity: in all structures chain B was contacting the aniline moiety of DRV whereas chain A was contacting the bis-THF moiety. (e.g. Fig. 4)

#### Molecular Dynamics Simulations

For each variant, three 100 ns simulations with randomized starting velocities were performed. The details of molecular dynamics simulation protocol have been described previously. ^58^ In short, forcefield parameters were assigned using the OPLS3 forcefield. ^55, 59^ The protein–inhibitor complex was solvated in a cubic box leaving at least 15 Angstrom between any solute atom and the periodic boundaries using the TIP3P water model. ^60^ Charges were neutralized by adding Na^+^ and Cl^-^, additional counterions were added up to concentration of 0.15 M. Thereafter the system was minimized in a series of steps. Simulations were run for 110 ns, where the first 10 ns equilibration periods were discarded for each simulation.

### Molecular Descriptors

Loss in inhibitor binding are a consequence of alterations in structure and in conformational dynamics. To quantify these changes and correlate them with the experimental enzyme inhibition, we calculated descriptors of protein-inhibitor and protein-protein interactions and dynamics. Python scripts to calculate the descriptors were developed in house and can be obtained from the accompanying Github repository. 3D coordinates from molecular dynamics trajectories were parsed using the Schrodinger Python api. The exact descriptors are detailed in the following section.

#### Van der Waals interactions

Pairwise van der Waals interactions between all protein residues and between protein and inhibitor residues were calculated from molecular dynamics simulations with 1 ns intervals. The OPLS3 forcefield parameters were used and calculations followed the standard combination and exclusion rules used with the OPLS force field. ^59, 61^

#### Hydrogen bonds

Inter- and intra-molecular hydrogen bond frequencies were calculated according to the geometric criteria defined in Steiner et al. ^62^ Water-mediated hydrogen bonds, that is hydrogen bonds between a solute atom, a solvent molecule and a third solute atom were also considered. All pairwise interactions were summed to residue-wise interactions.

#### Torsion angle entropy

To quantify torsion angle dynamics, a histogram of the ligand torsion angles and the f and Ψ angle distributions were constructed using a 36° bin width. The conformational entropy was calculated in terms of Shannon entropy. ^63^

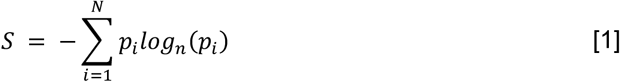

Here N=10 is the number of bins and p_i_ is the fraction of frames (from MD simulations) in which a torsion angle was in the configuration described by the i^th^ bin. The base of the logarithm defines the limits of the entropy distribution. Here the natural logarithm was used.

#### Root mean squared fluctuations

Root-mean-squared fluctuations (RMSF) of the protein C-alpha atoms, and root mean square deviations (RMSD) of the protein and inhibitor atoms relative to the starting structure were calculated.

#### Feature preprocessing

All calculated features were transformed to Z-scores (Equation 2) by subtracting the feature mean (µ) and dividing by the standard deviation (σ). Z-scores of the final test set and the cross-validation test sets were calculated using the training set mean and standard deviation, for each feature *x*:

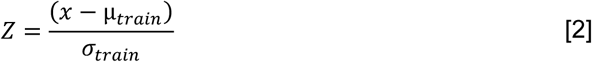

### Feature Selection

The objective of the feature selection protocol was to identify a sparse set of informative features, from which a model of the relationship between changes in physical interactions, dynamics and inhibitor binding can be established. Thus, we first identified the subset of features that are specific and informative.

#### Specificity

For each molecular feature that involved at least one variable protein residue, a regression model was constructed and compared with a competing null model. The null model was trained on a binary feature vector, with a value of 0 indicating an amino acid substitution relative to the NL4-3 wildtype. When the molecular feature involved 2 variable residues, the null model was trained on a n×2 feature matrix where n is datapoints. The models were compared using the Akaike information criterion (AIC).

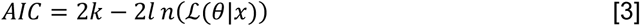

Here ℒ is the maximum likelihood value of the model given a set of parameters θ and k is the number of parameters. The relative likelihood of the competing models can thus be calculated using:

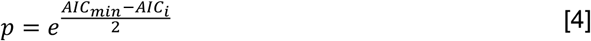

If the relative likelihood of the null model was below 0.05, the molecular feature was assumed to be specific. This means neither that molecular features that fail this test only encode for sequence information, nor does it mean that molecular features that are determined to be specific do not also encode for sequence information; however, the specific features capture more information than the sequence alone. The null model test was used only when the molecular features did not include the inhibitor.

#### Sparsity

The Scikit-Learn implementation of elastic net regression was used to reduce the high dimensional set of parameters obtained from molecular dynamics simulations. ^64, 65^. An L1-ratio of 0.75 and an alpha value of 1.0 were used. The elastic net model was trained on 66.6% of the training data. Training was repeated 100 times with different permutations of the training set. This was done to evaluate the stability of the coefficients. All non-zero (p<0.05, 1 sample t-test) coefficients were ranked according to their mean absolute values and a subset of parameters was chosen based on relative ranking (Fig. 2A)

### Regression Analysis

#### Model Selection

For the p selected features the relationship with the potency of the N points in the dataset was modeled using a linear model:

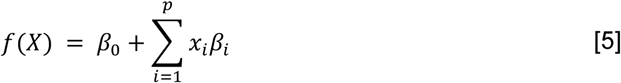

Where X is the N×p matrix of the selected features, βi is the coefficients of feature xi and β0 is the intercept. All possible combinations of the selected features were evaluated, and the models were compared using the Akaike information criterion. (Eq. 3 and Eq. 4). The statsmodels implementation of ordinary least squares regression and AIC were used. ^66^ All models with a relative likelihood of > 0.05 were considered equivalent to the best model and among these the model with the fewest parameters was chosen.

#### Model evaluation

Model performance on the training/validation set was evaluated using root-mean-square error (RMSE):

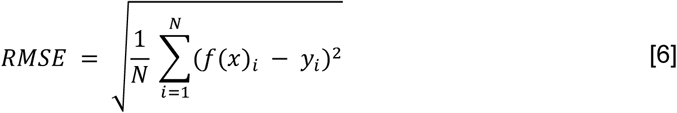

Pearson correlation:

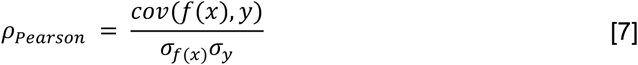

and Spearman correlation:

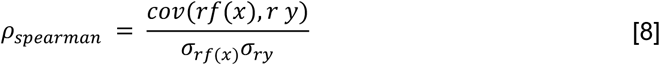

where N is the number of datapoints, f(x) is the prediction based on the independent variables x, y is the vector of dependent variables, cov(f(x), y) is the joined covariance, and σ_*f*(*x*)_σ_*y*_ are the standard deviations. In the case of Spearman correlation, the prefix “r” indicates that the continuous variables were transformed into ranks. In addition to the above metrics, the coefficient of determination (r^2^) was used to evaluate the goodness of a fit, where r^2^ is the square of ρ_pearson_ and represents the fraction of the variance in y explained by f(x).

### MM/GBSA Calculation

100 snapshots were extracted from MD simulations at 1 ns intervals. Previous studies of HIV-1 protease inhibitors using endpoint free energy calculations have employed similar sampling with simulation times between 4 and 60 ns simulations. ^35, 36, 67^ Once water molecules and counterions had been removed, MM/GBSA calculations were carried out using “*prime_mmgbsa*” as implemented in the Schrödinger 2018-1 release ^68^. Results were given from the single starting structure and averaged over the 100 snapshots from each trajectory. The reported mean and confidence intervals were calculated from 3 replicates.

## Acknowledgments

This research was supported by NIH P01 GM109767.

