## Supporting Information for "Deciphering complex mechanisms of resistance and loss of potency through coupled molecular dynamics and machine learning"

**Table S1**. Dataset information: HIV-1 protease variants with their experimental inhibition constants (K_i_) and PDB accession codes for available structures.

| **Name** | **K_i_ (nM)** | **ΔG (kcal/mol)** | **Set** | **Cluster** | **Reference** | **PDB ID** |
| --- | --- | --- | --- | --- | --- | --- |
| NL4-3 (WT) | 0.005 | -15.5 | Train | 2 | (51) | 6dgx |
| Var1-1Mut | 0.026 | -14.5 | Train | 2 | (51) | 6dh0 |
| Var2-2Mut | 0.23 | -13.2 | Train | 2 | (24) | 6opt |
| Var3-2Mut | 0.075 | -13.9 | Train | 2 | (51) | - |
| Var4-5Mut | 0.045 | -14.2 | Train | 2 | (19) | 4q1y |
| Var5-4Mut | 0.42 | -12.9 | Train | 2 | (24) | 6opu |
| Var6-8Mut | 12.8 | -10.8 | Train | 3 | (24) | 6opv |
| Var7-10Mut | 156.4 | -9.3 | Train | 3 | (24) | 6opy |
| Var8-11Mut | 759.2 | -8.4 | Train | 3 | (24) | 6opz |
| Var9-9Mut | 77.6 | -9.8 | Train | 3 | (50) | - |
| Var10-7Mut | 12.1 | -10.9 | Train | 3 | (50) | - |
| Var11-10Mut | 57.6 | -9.9 | Train | 3 | (50) | - |
| Var12-6Mut | 3.9 | -11.5 | Train | 3 | (50) | - |
| Var13-10Mut | 26.0 | -10.4 | Train | 3 | (50) | - |
| Var14-12Mut | 12.1 | -10.9 | Train | 3 | (50) | - |
| Var15-12Mut | 58.3 | -9.9 | Train | 3 | (50) | - |
| Var16-10Mut | 57.0 | -9.9 | Train | 3 | (50) | - |
| Var17-14Mut | 11.7 | -10.9 | Train | 3 | (50) | - |
| Var18-20Mut | 0.75 | -12.5 | Test | 1 | (69) | 3ttp |
| Var19-20Mut | 15.0 | -10.7 | Test | 1 | (70) | 3u7s |
| Var20-7Mut | 0.026 | -14.5 | Test | 2 | (71) | 3ekt |
| Var21-24Mut | 6.95 | -11.2 | Test | 1 | (13) | - |
| Var22-4Mut (SF2) | 0.014 | -14.9 | Test | 2 | (72) | 1t3r |
| Var23-6Mut | 6.6 | -11.2 | Test | 2 | (73) | 2f80 |
| Var24-6Mut | 17.0 | -10.7 | Test | 2 | (74) | 3cyw |
| Var25-20Mut | 31.0 | -10.3 | Test | 1 | (75) | 3ucb |
| Var26-6Mut | 0.51 | -12.8 | Test | 2 | (76) | 5kqy |
| Var27-6Mut | 1.6 | -12.1 | Test | 2 | (74) | 3d1z |

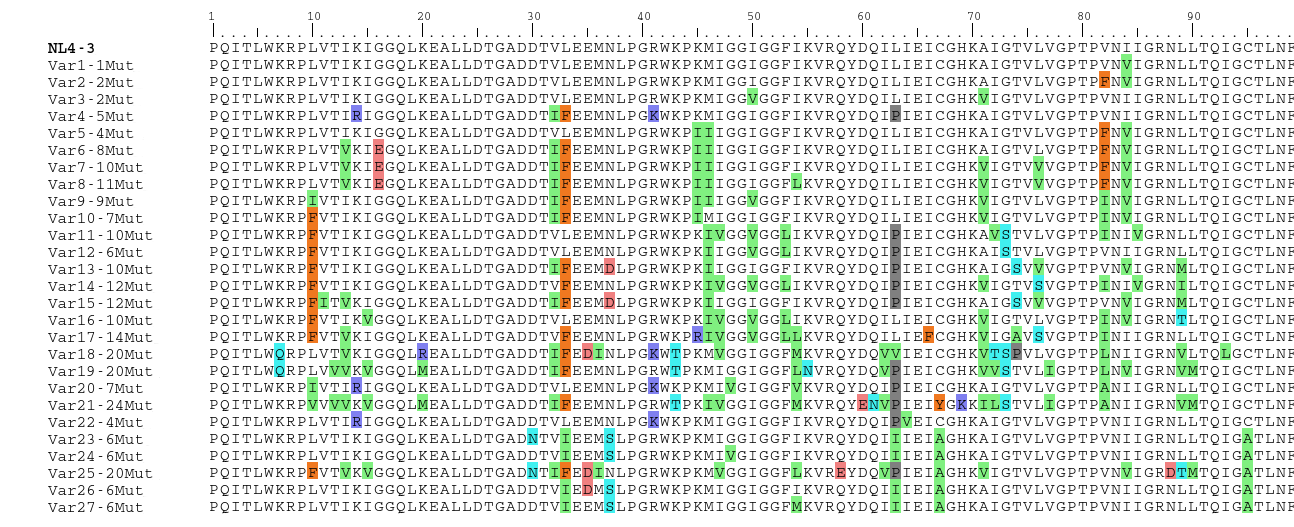

**Figure S1.** Amino acid sequence alignment of HIV-1 protease variants in the dataset. Mutations relative to the reference wild type (NL4-3) are colored according to residue type. (Green: Hydrophobic; Orange: Aromatic, Red: Negative Charge; Blue: Positive Charge; Cyan: Polar; Gray: Small)
